# Performance of Naiive Spectral Geometric Models in Histopathology AI

**DOI:** 10.64898/2026.01.30.702908

**Authors:** Alejandro Leyva, M. Khalid Khan Niazi

## Abstract

There have been no systematic evaluations of purely spectral models for digital pathology tasks. We implemented and benchmarked four pipelines: binary classification on the BreaKHis dataset, multi-class region classification in glioblastoma, spatial transcriptomics, and denoising on Visium 10x. Across all tasks, extensive cross-validation and grouped splits showed that purely spectral models did not improve performance over CNN-only baselines, but offer useful complementary tools for interpretability and processing. Denoising showed strong performance that proves utility in data-scarce or heterogeneous image environments. Equivalence testing confirms that spectral and CNN model performances fall outside ±3% AUC. Fusion models between CNNs and spectral models show higher balanced accuracy. Spectral models failed to generalize across spatial transcriptomics tasks, with low correlation despite stable training loss. These findings represent a systematic negative result: despite their theoretical richness, spectral geometric features and SNO embeddings prove to be complementary features for WSI classification or segmentation. Reporting such outcomes is essential to establish empirical boundaries for spectral methods and to encourage future work on conditions or data modalities where these approaches may hold greater promise.

## 1 Introduction

Spectral mathematics seeks to describe the behavior of eigenvalue distributions across matrices and higher-dimensional spaces (1, 2). In histopathology, H E staining followed by microscopic examination can be interpreted through geometric variations that spectral methods quantify. Each structure is distinguishable by its curvature and topology, reflecting underlying cellular mechanisms (3). Spectral imaging has been applied to histopathological analysis since the 1990s, motivated by the ability to simultaneously observe multiple molecular signals within the same tissue (4). Although hyperspectral methods offer molecular detail, hyperspectral imaging has yet to consistently outperform vision transformer (ViT) foundation models for histopathology tasks, primarily due to data limitations (5). We instead focus on spectral geometry and Laplace-derived operators on tissue structure. Spectral theory has contributed significantly to AI, most notably in improving image quality, denoising, and signal separation (6). Pilot studies have shown that combining spectral imaging with convolutional models can improve diagnostic accuracy (7, 8, 9).

While deep learning methods have been widely applied to histopathological images, there has been no systematic evaluation of purely spectral geometric deep learning for these tasks. We hypothesized that spectral geometric techniques could extract interpretable features inaccessible to traditional models, and that these features could improve accuracy in both hybrid and standalone models. To test this, we configured several models to extract spectral features, which were then processed through conventional deep learning methods, as well as through families of spectral neural networks. Each approach was evaluated across four representative tasks: classification, segmentation, spatial transcriptomics, and denoising. Spectral geometry provides a natural framework for this investigation. By encoding curvature, topology, and morphodynamics into sets of spatially distributed eigenvalues, we obtain features that reflect the functions necessary to generate specific morphologies. Higher-index eigenfunctions correspond to higher-resolution representations of shapes (10). The overarching question is whether spectral geometric methods can capture morphologically discriminative information in histopathological images. Each task was chosen to test a specific property: the ability to globally discriminate between images and patches, locally resolve features within patches, predict feature distributions within a region, and refine and denoise features at high resolution.

Unlike CNNs or ViTs, spectral approaches are directly interpretable: they operate on precomputed eigenvalues that encode curvature, shape, and topology (12, 13). We concatenated spectral features derived from multiple methods into unified feature vectors, which were then input into CNNs or spectral neural networks (SNNs). SNNs, originally developed in particle physics, learn solutions to partial differential equations by operating in the frequency domain before mapping results back to the spatial domain (14, 15). JacobiConv, for instance, projects features into a spectral graph basis and learns flexible weights for Jacobi polynomial expansions. Spectral neural operators (SNOs) instead transform features into the frequency domain and process them with nonlinear operator layers. These architectures offer a unique ability to capture nonlinear feature interactions that remain invisible in the spatial domain.

## 2 Materials and Methods

Several mathematical techniques were employed on the resulting eigenvalue matrices. Diffusion models, for example, were used to characterize morphology distributions around specific points (11). Spectral techniques have already shown promise in pathology: elliptical Fourier descriptors, for example, achieved 88 percent accuracy in a convolutional model for histology detection (16). Building on this, our study explores whether spectral- or frequency-domain derived features are more effective when paired with spectral deep learning models. We applied multiple methods, including Heat Kernel Signatures (measuring differences in contour diffusion over time), fractal encodings (quantifying lacunarity), graph wave energy (capturing local transitions at boundaries), Slepian functions (cross-analyzing curvature-defined regions), and wave kernel analysis (estimating prominence probabilities across points). Together, these methods probe the extent to which spectral mathematics can provide new, morphologically meaningful features for histopathological imaging. A fixed grid of 64×64 was constructed across each 512×512 image patch. Nodes were defined by the proportion of tissue pixels within each grid cell, with a tissue threshold of 20 percent. All computations were performed using Ohio State Supercluster’s Ascend desktop (17) on NVIDIA100 GPUs on Cuda 11.8.0. For each valid node, eight features were computed: mean grayscale intensity, local entropy, Laplacian variance, mean hematoxylin and eosin channel intensities, saturation, Sobel gradient variance, and tissue-to-boundary ratio. Each node was connected to its four immediate neighbors, forming a local mesh. Edge weights were computed using Gaussian weighting to preserve smooth transitions between adjacent nodes.

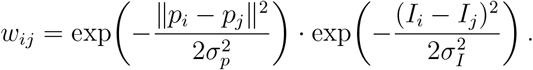

The normalized Laplacian was then computed as:

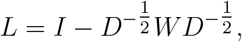

where *W* is the adjacency matrix, *D* is the diagonal degree matrix, and *I* is the identity matrix. After computing the Laplacian, eigenvalues *λ*_*k*_ and eigenvectors *u*_*k*_ were obtained by solving the eigenvalue problem:

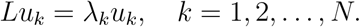

The spectral coefficents were computed by applying the 8 channel features extracted onto the eigenvalues for each node, where X refers to the channel vectors, and a refers to the spectral coefficients, and U is the solution to the eigenvalue problem:

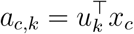

From these eigenpairs, multiple spectral features were extracted:

— Heat Kernel Signature (HKS):

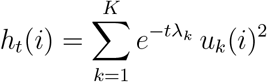
— Spectral Wavelet Energy:

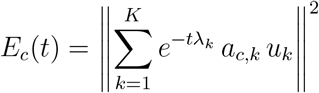
— Slepian Concentration:

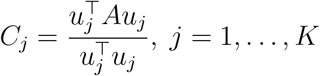
— Fractal descriptors: log–log slopes, Rényi dimensions, lacunarity

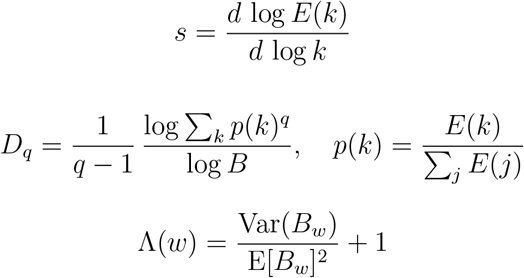
— Persistent homology: lifetimes and persistence entropy.

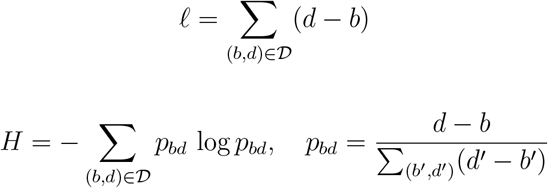

The signature coefficients of the heat kernel described as lambda were between 0 ¡0.02 and were concatenated onto the original channel vector to measure the distance between the curvatures of each tissue block within a selected grid. The spectrum wavelet energy features extract the magnitude of the distances across the spectral coefficients labeled a, creating a localized measurement of distance around each instance. Slepian concentrations were observed by using the adjacency matrix A to ensure that the grids that were filtered would not pass through the sequence, and u refers to the jth eigenvector within the eigendecomposed eigenvector generated by solving the eigenvalue problem using the eigensh() function. This measures the local energies of nodes, whereby higher entropy suggests less cohesive cellular or morphological boundaries, and serves to differentiate spatial regions by their eigenvalues. Within the fractal embeddings, the log-log scale was computed by taking the normalized entropy of each spectral coefficient with respect to channel c and frequency band k, measuring the rate of decay across each node boundary. Renyi dimensions computed were within channels 0, 1, 2, where channel 1 is reduced to shannon entropy, 2 contributes to correlation between clusters, and channel 3 is correlation across high energy bands B, where eigenvalues of certain magnitudes (frequency intervals) are placed into cluster B. p(k) is calculated by taking energy E of a particular band and calculating the entropy across all bands. This was useful for measuring cross-correlational insights across the image, and measuring the ordered states within the tissue. Lacunarity was used to measure the heterogeneity of patterning across the image by computing the energy across spectral coefficients with the summations of energy within band window B and computing the entropy relative to the mean of B. The increase in variance of band B quantifies the heterogeneity of the patterning within certain nodes. Persistent homology was used to measure the strength of various motifs in the image, including loops, cuts, and whitespaces.Gudhi was used to compute the birth and death (b,d) of the features expanding from the node at radius epsilon. First, a graph is constructed using the nodes and spectra coefficients computed from the laplace-beltrami decomposition. If the pairwise distance is established between all points p,q, then each set of edges will continue to expand in a simplex as long as d(p,q) ¡ epsilon. as epsilon expands, features of the original point will disspiate and the edges between each node will be pruned. The number of births counts the number of nodes formed between nodes at scale b, and vice versa. The vietoris-Ris complex is described, whereby set of nodes forms a simplex as radius epsilon expands:

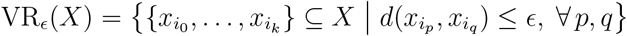

As radius epsilon expands, more simplex formations (triangles, tetrahedrons) are formed:

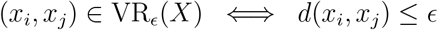

As the radius expands, edges and nodes are pruned due to the dissipation of the motif via thresholding and component merges:

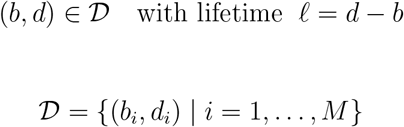

The rate of change of (b,d) is used to compute the persistent entropy across the tissue sample, which can then be used for the persistent homology formulas described below.

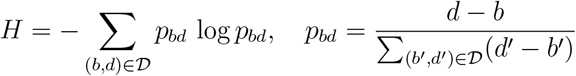

After feature extraction, the vectors are concatenated into a 1D vector of about 180 features. The various windows, channels, and k values are all described in the Python script. The vector is reduced to 25 dimensions using principal component analysis (PCA).

**Table 1:** Architecture of the Spectral Neural Operator head.

| Layer | Operation / Params | Tensor shape |
| --- | --- | --- |
| Input | — | $(B, C, K)$ |
| SpectralMix 1 | $U \mapsto \phi(B[UA] + \beta)$; $A \in \mathbb{R}^{K \times K}$ (low-rank $r$), $B \in \mathbb{R}^{C \times C}$, $\phi = \text{Softplus}$ | $(B, C, K)$ |
| Dropout 1 | $p = 0.10$ | $(B, C, K)$ |
| SpectralMix 2 | same as above (shared hyperparams, new weights) | $(B, C, K)$ |
| AdaptiveAvgPool<sub>K</sub> | mean over spectral axis $K \rightarrow 1$ | $(B, C, 1)$ |
| Flatten | squeeze last dim | $(B, C)$ |
| Linear 1 | $C \rightarrow H$ | $(B, H)$ |
| Softplus | — | $(B, H)$ |
| Dropout 2 | $p = 0.10$ | $(B, H)$ |
| Linear 2 (head) | $H \rightarrow n_{\text{cls}}$ | $(B, n_{\text{cls}})$ |

*Aux output (for fusion): pooled embedding $z$ taken after AdaptiveAvgPool — $(B, C)$ .*

### A. Segmentation

The GBM-WSSM glioblastoma dataset (n=83) was used to train, test, and validate the segmentation model using a LOSO strategy employing cross-kfold validation. Segmentations were pre-annotated as to provide a ground truth for the model. The model was told to predict regions of pseudopalisading necrosis, necrosis, infiltrating tumour, The model was compared against a CNN using LightGBM, SNO, SNO and the LGBM, and L2 regression. All images were at the same magnification.

### B. Classification

The Breakhis breast cancer dataset composed of patches (n=7258) was used to test,train, and validate the model using the same training configurations as shown in figure 1. The model was tasked to classify between benign and malignant tumours. the dataset is split into 5x,10x,20x, and 40x magnifications.

**Figure 1:**
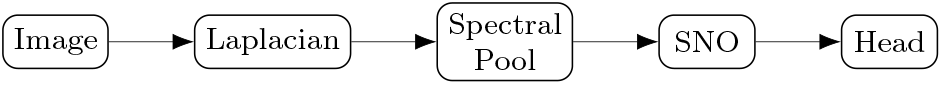
Spectral pipeline: Image → Laplacian → Spectral Pool → SNO → Head.

**Figure 2:**
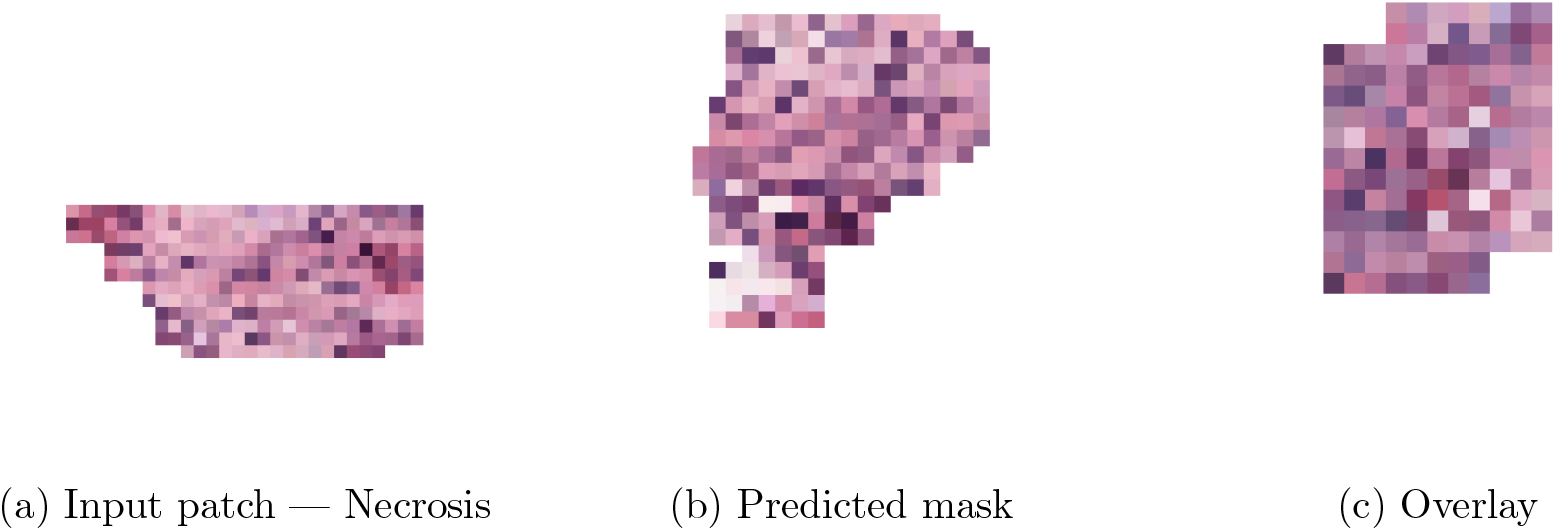
Qualitative segmentation results from the spectral model on necrotic and perinecrotic regions. The model identifies necrotic tissue boundaries and viable zones consistent with histological annotation.

### C. Spatial Transcriptomics

The Visium 10x breast cancer dataset (n=42) was used to train, test, and validate the model. Metrics of accuracy for this model include Pearson’s correlation coefficient (PCC), and Mean squared error (MSE). Genetic prediction was validated using crossfold validation and LOSO to prevent leakage. D. Denoising: The Visium dataset for breast cancer was used, though this time the model was trained spot-wise rather than across an entire image to produce synthetic spots and improve the accuracy of a baseline model. The SNO was compared against a kNN and ralphensemble model. Laplacian smoothing was performed and synthetic spots were created by predicting gene expression within the spot.

## 3 Results

The results for segmentations show as follows, where FULL refers to all features passing through the SNO:

**Table 2:** GBM Global Classification (averaged across folds)

| Model | Accuracy ↑ | Balanced Acc ↑ | mAUC ↑ |
| --- | --- | --- | --- |
| SNO (Spectral) | 0.26 | 0.22 | 0.61 |
| CNN → LGBM | **0.54** | **0.41** | **0.80** |
| FULL → LGBM | <u>0.43</u> | <u>0.31</u> | <u>0.66</u> |
| FULL → LR | 0.34 | 0.27 | 0.65 |

GBM global segmentation suffered from class imbalance, as certain classes such as perinecrotic zones were not evenly dispersed within the dataset, leading to sparse data to train the models to detect these classes. While the low performance of the Convolutional neural network is due to the dataset, the purely spectral model does not perform well on classification tasks. The spectral neural operator trained alone on spectral features performed worse than a standard lasso regression, naturally because the model fails to discriminate when using non-spectral features. Light gradient boosting (LGBM) increased the mAUC by 5% within the FULL spectral architecture. For the classification task, spectral neural operators prove to be unoptimized for segmentation and are not yet suited for generalized embeddings. However, the purely spectral models that use both spectral features and spectral operators prove to capture a decent, competitive signal. The results for the classification task, where fused relies on the outer product within the features. the spectral and convolutional features were reduced using PCA such that the features together could be tested with a 50/50 split.

**Table 3:** BreakHis Global Classification (5-fold averages)

| Model | Accuracy ↑ | Balanced Acc ↑ | AUC ↑ |
| --- | --- | --- | --- |
| SNO (baseline) | $0.65 \pm 0.21$ | $0.52 \pm 0.05$ | $0.62 \pm 0.12$ |
| SNO → LGBM | $0.73 \pm 0.05$ | $0.56 \pm 0.04$ | $0.59 \pm 0.11$ |
| CNN → LGBM | **$0.83 \pm 0.06$** | $0.73 \pm 0.10$ | **$0.80 \pm 0.13$** |
| FUSED (SNO $\oplus$ CNN → LGBM) | <u>$0.82 \pm 0.06$</u> | **$0.73 \pm 0.08$** | <u>$0.78 \pm 0.13$</u> |

**Table 4:**
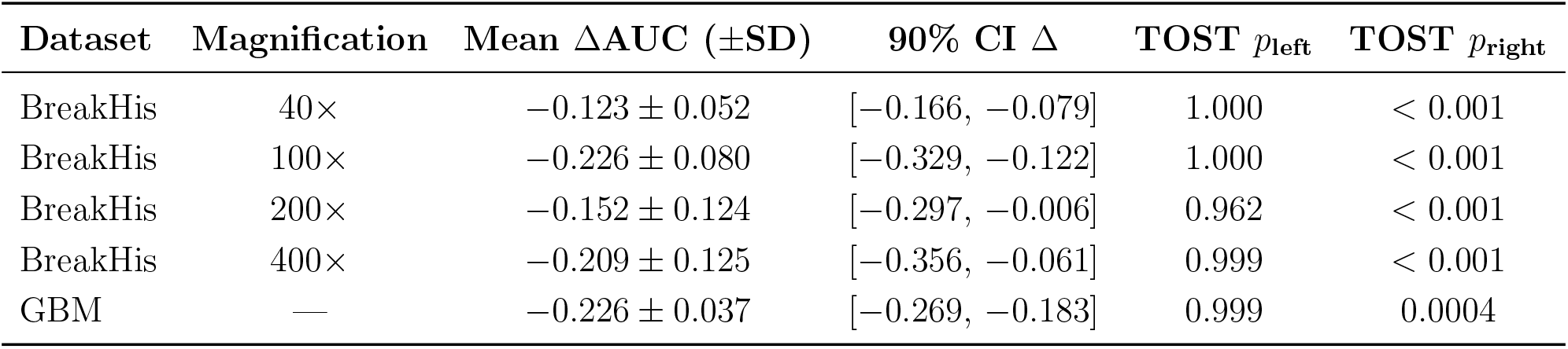
Equivalence testing (TOST) results comparing Spectral vs. CNN models. ΔAUC = AUC(Spectral) − AUC(CNN); equivalence margin Δ_0_ = 0.03, *α* = 0.05.

The results demonstrate that purely spectral models are not competitive with current methods. Neither the spectral architecture or the full spectral features and architecture could compete within a 10% AUC range. Interestingly, when these spectral signals were fused with Convolutional features, the balanced accuracy increased, while the AUC only slightly decreased. For both the pure CNN and Fused models, the vector size was the same in both conditions in order to create a fair comparison between the Fused and CNN models. however, the fused model adds the spectral features, and shows a decrease in AUC due to invariant features such as Slepian embeddings, but demonstrates a higher balanced accuracy. Spectral features demonstrate and ability to provide complimentary interpretable features that can aid the performance of CNNs. The conclusion between the two tasks is that purely spectral models can not outperform CNNs, as the performance discrepancy between the models across all magnifications and tasks prove to be statistically significant using a twosided T-test (TOST) with in a 90% confidence interval (CI). However, the scope is limited by the development of spectral architectures towards these specific tasks. In the BreakHis classification task, spectral features prove to be complimentary to convolutional features, and can perhaps provide interpretability without damaging performance.

Cross validation learning curves in figure 3 show that the learning trends for the breakhis classification task for each fold across magnifications. Similiar learning trends are demonstrated across all models, though performance worsens when the 400x magnification is reached. The pure CNN model has the best learning rate across all models, though the fused model sometimes outperforms the CNN in Fold 4 of the 100x magnification, fold 2 of 200x, and fold 3 of 400x. Fold 5 of 100x was excluded due to parsing and class imbalance errors. The comparisons between the purely spectral architecture (SNO) and the CNN show that the SNO model fails to generalize across magnifications. However, there are occasional cases by which SNO outperforms CNN, demonstrating that the features analyzed by the model present a meaningful signal. The GBM dataset experienced severe class imbalance on fold 3 and was excluded, as certain sparse classes simply weren’t present. Regardless, the comparison holds to the dimensionality reduction and comparison design. The CNN simply adapts to class imbalance with better resilience than spectral models. The results for the spatial transcriptomics database using JacobiConv: For the spatial transcriptomics task, performance on breast cancer on visium 10x (n=40) was taken from (citation), using a cross fold LOSO architecture, demonstrated very poor performance against traditional models. Training spot-wise, interestingly, demonstrated excellent performance for the denoising task shown in figure 5, which uses spectral mappings to smooth images and create mini-spots, or increase resolution around patches, and then predict the gene expression at that position. The distribution of predictions per case demonstrates high MSE and MAE with very low PCC, with the highest count of RMSE being .35, .07 for PCC, .22 for MAE. When simulated with data scarcity (only 25% of spots available) across multiple shuffle runs, spectral mappings on average surpass kNNs and RBFs for denoising. Across Normal conditions, kNN and RBF outperform spectral mappings (SSR) for spot-based predictions. Spectral mappings perform well in sparse data conditions and provide an excellent tool to map within-slide heterogeneity, but the features extrapolated can not be generalizable across cases. Spectral geometric models prove to be complimentary features that can increase the resolution of hetereogenous structures within data-scarce partitions. Figure 6 shows the performance of SSR within non data scarce environments, where both kNN and RBF outperform SSR across all metrics. Spectral mappings in non data scarce environments did not improve gene prediction performance for other models, performance decreased in figures (e-h).

**Table 5:**
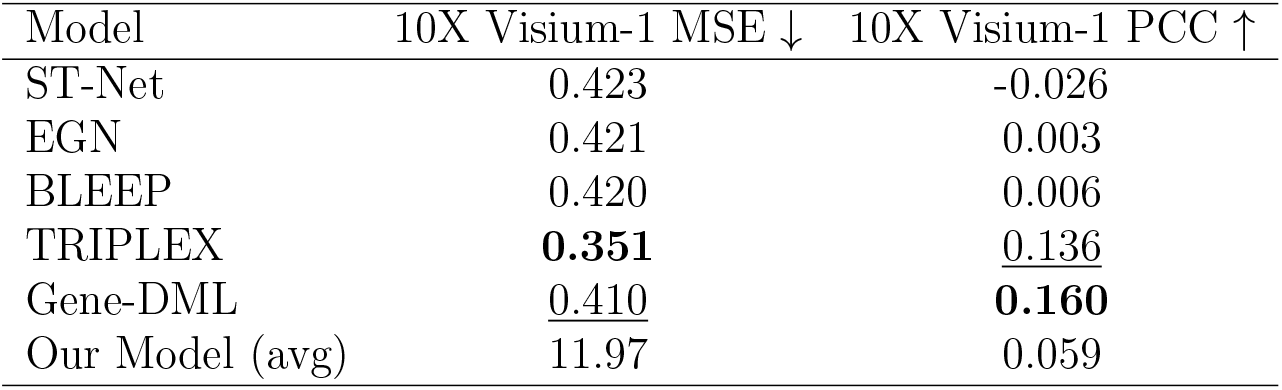
Spatial transcriptomics prediction performance on 10X Visium-1.

**Table 6:** Performance comparison across smoothing methods for spectral ST prediction. Higher PCC and *R*^2^ indicate better performance; lower RMSE and MAE are better.

| Method | PCC | MAE | RMSE | $R^2$ |
| --- | --- | --- | --- | --- |
| kNN<sub>xy</sub> | 0.0959 | 0.2478 | 0.3415 | -0.074 |
| RBF<sub>xy</sub> | 0.0707 | 0.2472 | 0.4152 | -0.560 |
| SSR (ours) | 0.1760 | 0.2421 | 0.3298 | -0.015 |

**Figure 3:**
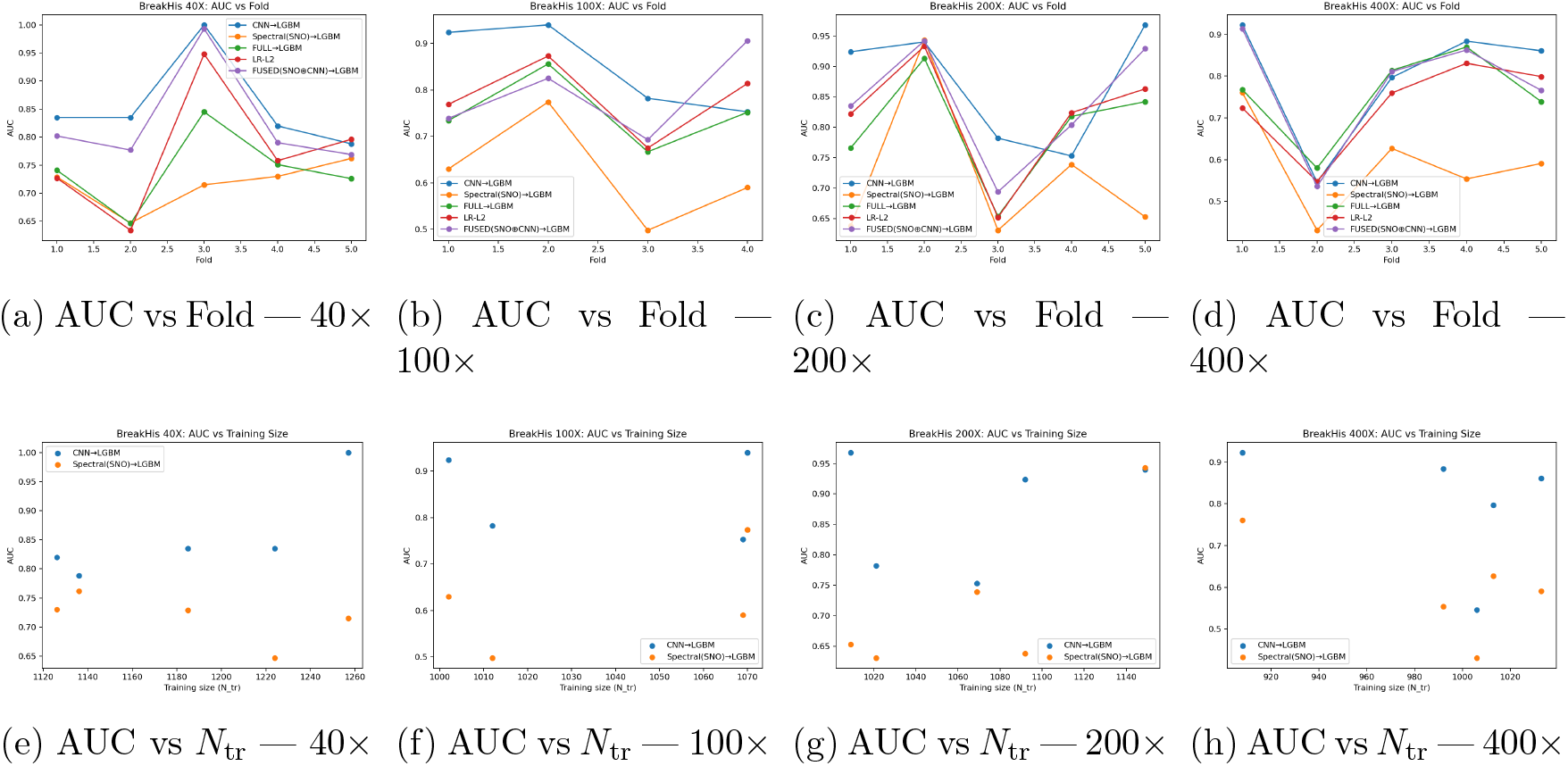
Cross-validation learning curves on BreakHis across magnifications. (**Top**) AUC vs fold for each magnification (CNN→LGBM, Spectral(SNO)→LGBM, and others). (**Bottom**) AUC vs training set size *N*_tr_ for the same magnifications.

**Figure 4:**
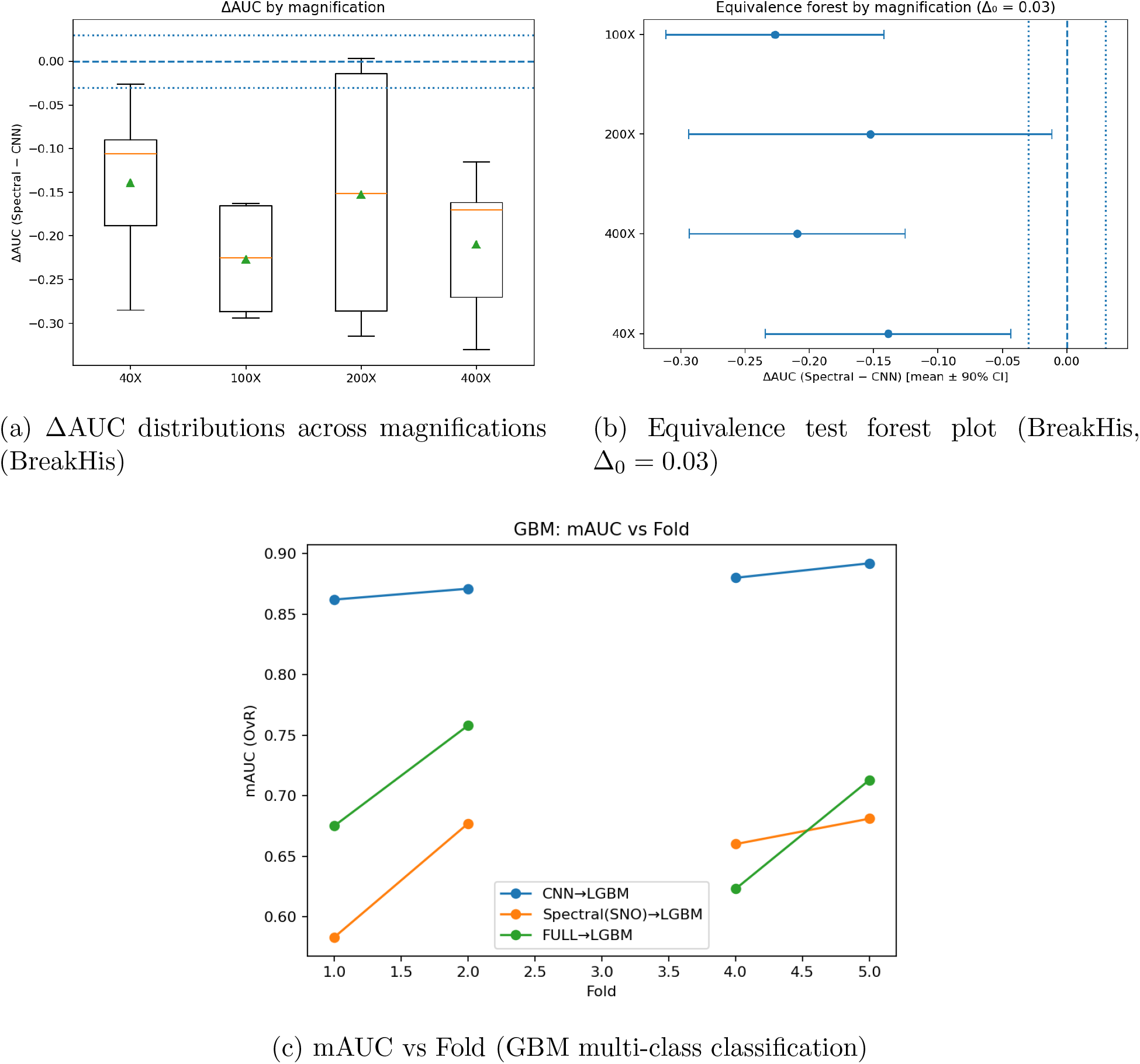
Summary of cross-validation equivalence analyses. (**Top left**) Distribution of ΔAUC values across BreakHis magnifications. (**Top right**) Forest plot showing mean ΔAUC *±*90% CI with equivalence bounds (Δ_0_ = 0.03). (**Bottom**) GBM learning curve showing mAUC vs fold for CNN→LGBM and Spectral(SNO)→LGBM models.

**Figure 5:**
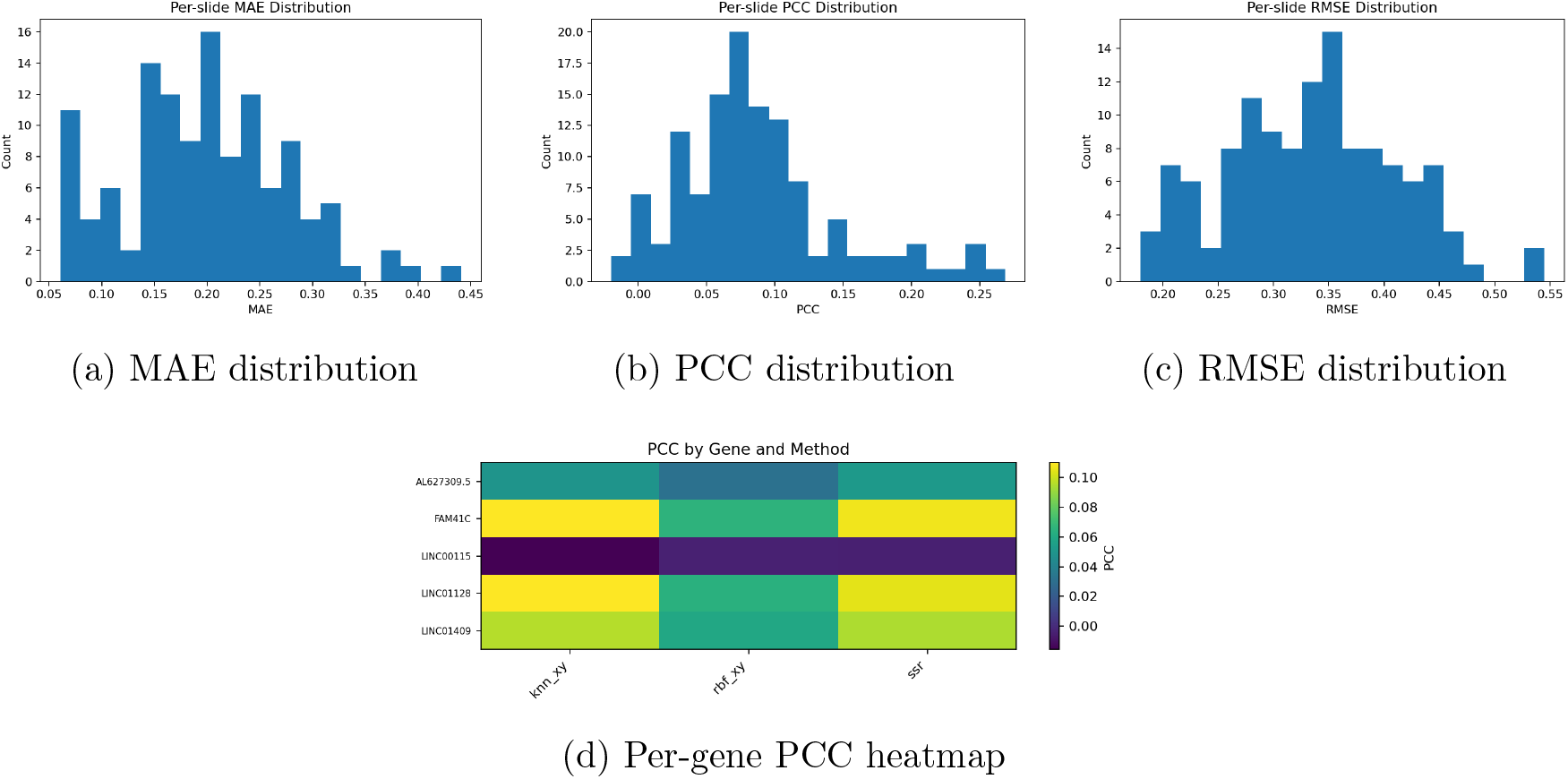
Distribution of regression metrics across genes and samples. (**Top**) Histograms of mean absolute error (MAE), Pearson correlation coefficient (PCC), and root mean square error (RMSE) across all predictions. (**Bottom**) Gene-wise correlation heatmap illustrating PCC variation across the dataset.

**Figure 6:**
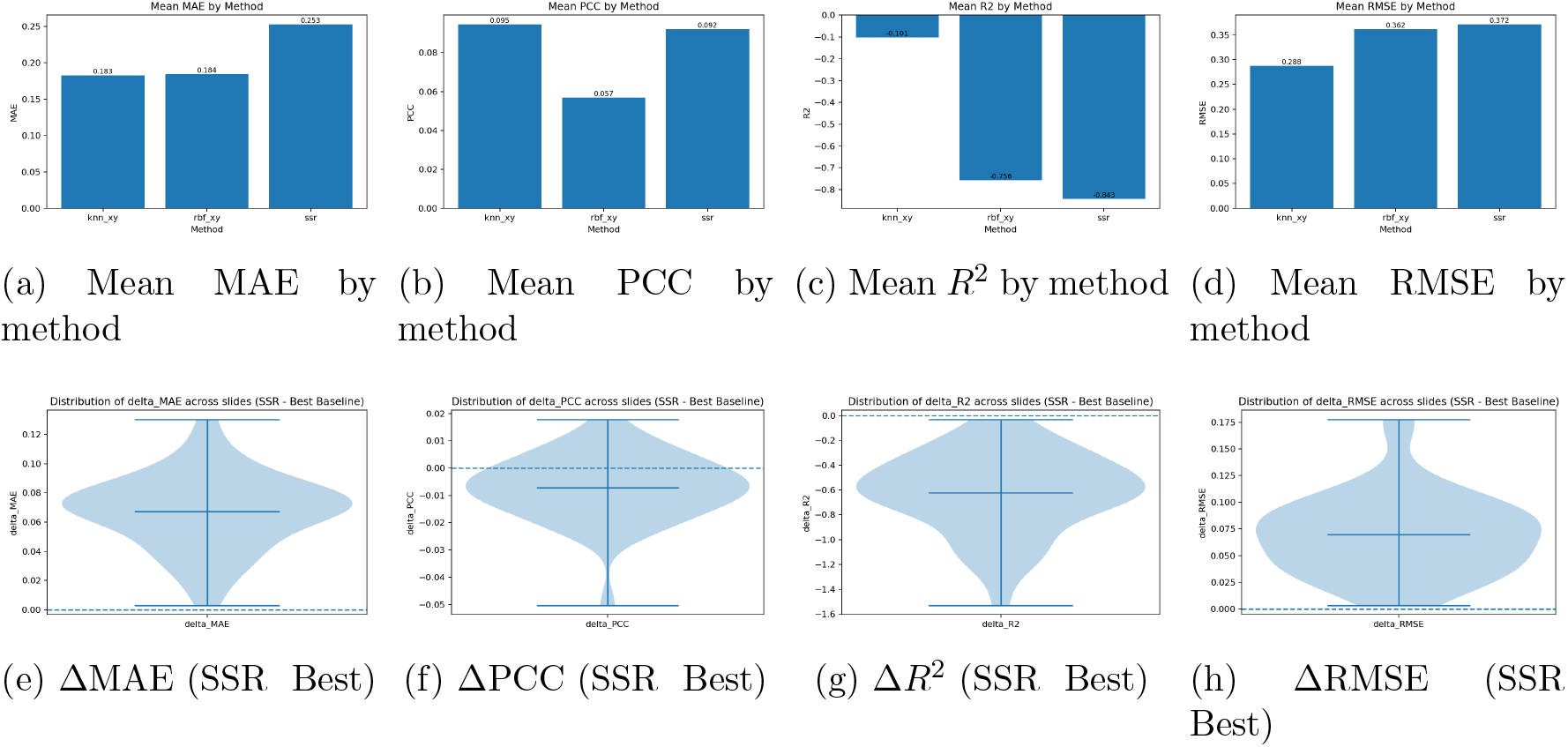
Comparison of regression metrics across methods. (**Top**) Mean absolute error (MAE), Pearson correlation (PCC), coefficient of determination (*R*^2^), and root mean square error (RMSE) averaged across models. (**Bottom**) Violin plots showing the distribution of metric deltas (Δ = SSR Best) across folds. Negative values indicate performance gains from the best model relative to SSR.

The results for the denoising tasks, such that the average accuracy across each user was compared.

As shown in figure 7, the training curves for the spectral spatial transcriptomics model were relatively stable and presented poor results for gene correlation. validation loss was relatively improved, though suboptimal. Figure 8 shows that the graphical construction for laplace beltrami transformation are aided by the hematoxylin staining by highest priority, which provides an understanding of what structures spectral networks gravitate towards to visualize geometric features. Spectral features do extract relevant geometric signal from images, as staining is the means by which larger models such as Vision Transformers implicitly spatially autocorrelate between pixels within H & E images.

**Figure 7:**
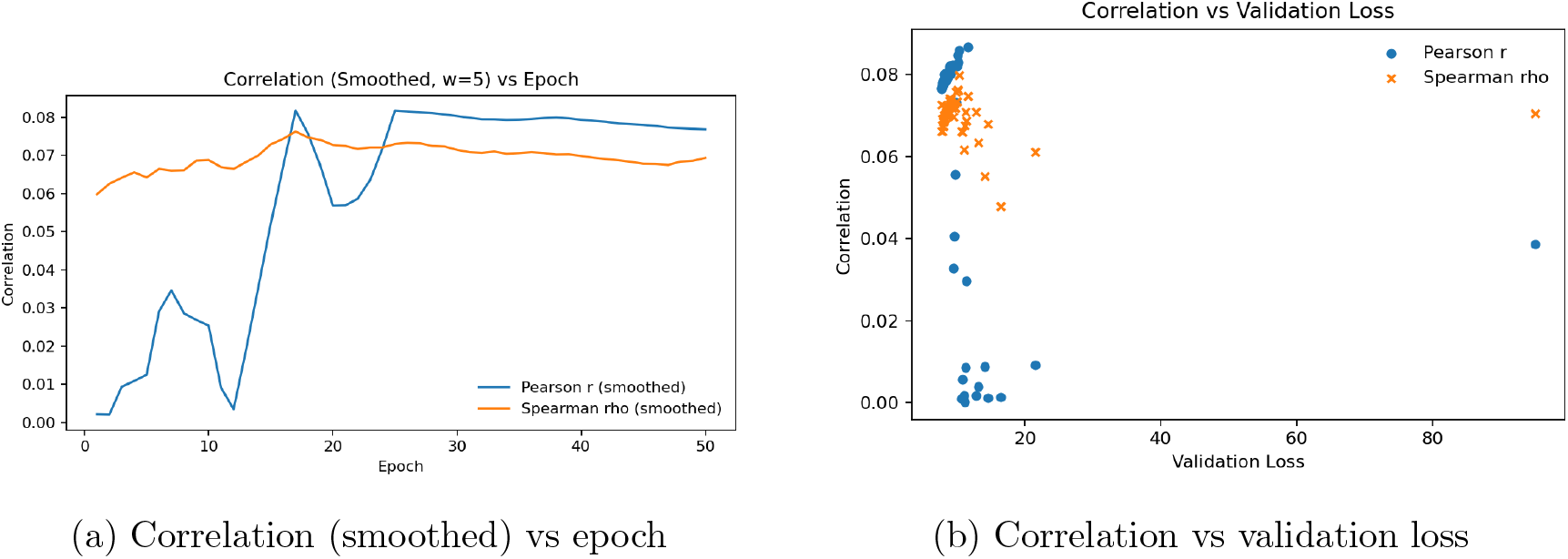
Spectral ST prediction learning behavior. (A) Pearson’s *r* and Spearman’s *ρ* across epochs (rolling mean *w*=5). (B) Correlation–loss relationship showing marginal signal recovery under low validation loss.

**Figure 8:**
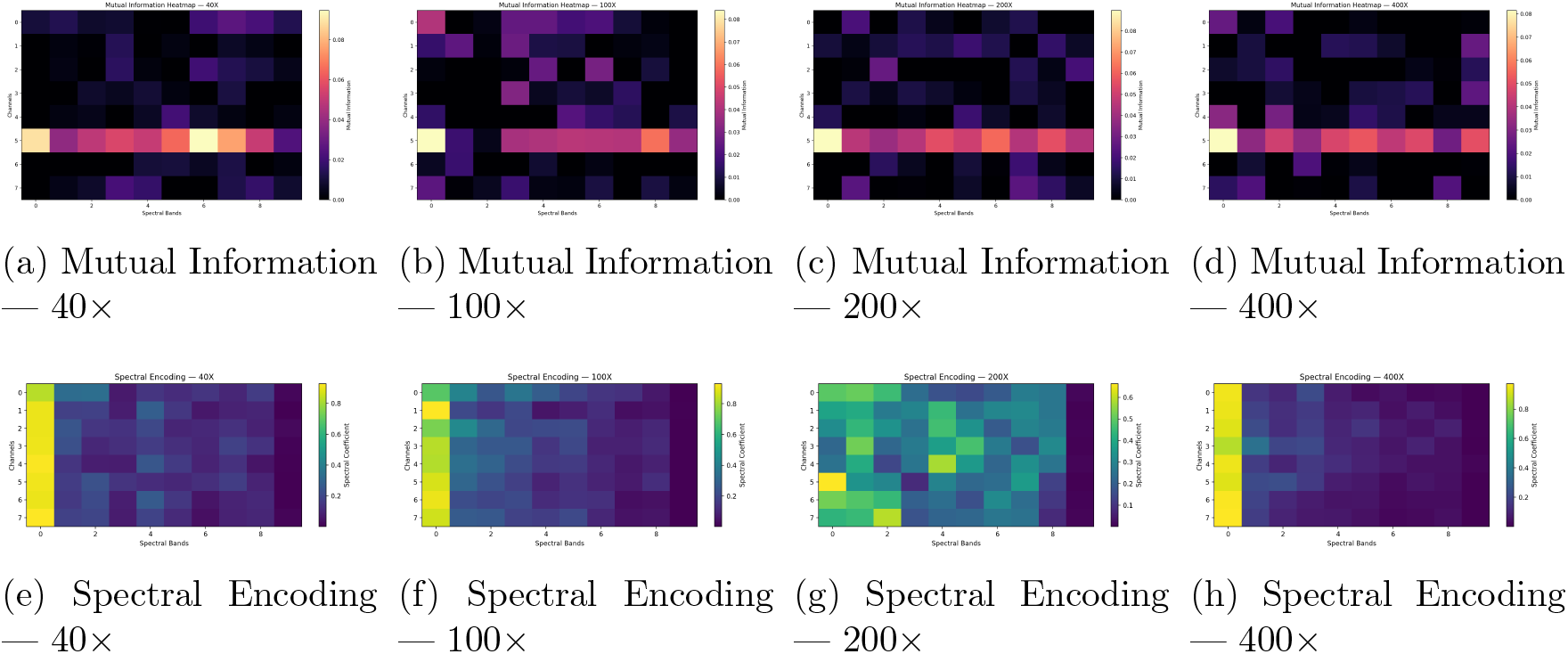
Spectral analysis of BreakHis histopathology patches across magnifications. (**Top**) Mutual information between spectral bands and RGB channels showing concentration of inter-band dependencies around channel 5. (**Bottom**) Spectral encoding coefficients illustrating how spectral energy distribution broadens with magnification.

## 4 Discussion

Overall, the spectral pipelines produced stable but consistently lower performance than CNN baselines across classification, segmentation, and ST prediction. Balanced accuracy improved in fused models, confirming that spectral features carry complementary signal. Spatial transcriptomics performed poorly in cross-slide settings but improved substantially in spot-wise denoising under sparse conditions. These findings establish both the boundaries and concrete utility cases of spectral geometry in pathology. In practice, spectral extraction and training took 45 minutes per epoch on an A100 GPU. While spectral features capture meaningful geometric and textural information, they remain weaker than conventional convolutional methods. Even with large training datasets, the spectral model struggled to generalize across independent images. The relatively strong within-slide performance suggests that spectral representations can predict global morphological patterns relevant to cancer subtyping; however, their inclusion alongside convolutional features did not improve overall accuracy.

In the segmentation task, the model was hindered by class imbalance and primarily detected regions with high class prevalence. Because spectral features are intrinsic to each image, the model cannot easily transfer information learned from previously computed features—particularly evident in the spatial transcriptomics task, where high-dimensional gene expression could be predicted at the spot level but not the slide level. Nevertheless, the spectral framework retains a notable advantage in denoising and resolution enhancement for small datasets, where localized information dominates and data scarcity limits end-to-end convolutional approaches.

Although JacobiConv achieved modest gains for spatial transcriptomics and denoising, the Spectral Neural Operator (SNO) still showed promise for processing spectral representations efficiently. Interestingly, the learned spectral coefficients correlated most strongly with the staining intensity across the eight input channels, suggesting that the morphology detected by the model is primarily outlined by staining boundaries rather than complex tissue geometry. In spatial transcriptomics, spot-level models using low-frequency Heat Kernel Signature (HKS) channels exhibited strong spatial autocorrelation, yet leave-one-slide-out validation revealed that these localized contour distances could not be generalized across samples.

Ablating the Slepian function features reduced classification performance by approximately 20 percent, indicating that these features contribute nontrivially to the overall representation. However, expanding to higher frequency bands produced largely invariant signals, implying that localized spectral coefficients rarely exhibit high-frequency variation, even in tumor-dense regions. Persistent homology detected recurring motifs within small neighborhoods, showing potential for localized pattern recognition.

Earlier iterations of the model also included Wave Kernel Signatures and Geometric Fourier transforms, but these features failed to carry generalizable signal. The probability of a feature occurring at a specific point proved too patch-dependent to support robust learning. Similarly, graph wavelets highlighted local distinctions in segmentation and classification but did not exhibit spatial autocorrelation in gene prediction tasks. Fractal features mirrored the performance of Slepian descriptors, contributing minimally to overall model strength. Because the current approach is patch-based, fractal networks cannot be easily embedded within graph structures. Previous fractal convolutional architectures have shown promise in histopathology AI, but graph networks do not yet provide spatially consistent frameworks for fractal encoding.

For future work, balancing localized and globalized spectral representations will be essential. Features that are too local fail to generalize, while those that are too global lose morphological specificity. The present model demonstrates potential for image smoothing and resolution enhancement in spatial-omics applications, yet its ability to generalize across slides remains limited. Frequency bands can be fine-tuned for individual tasks, but optimal settings vary widely by staining type and coefficient normalization.

A more fundamental constraint arises from reliance on the normalized Laplacian. Alternative formulations—such as Schrödinger or PDE-based Laplacians—yield eigenvalues that are numerically unstable under tensor computation, limiting practical experimentation. Nonetheless, many spectral operators remain unexplored in the context of histopathology AI. With a clearer understanding of the specific geometric and morphological features these methods capture, future work can integrate them into interpretable, hybrid models that bridge mathematical elegance with biological relevance.

## 5 Limitations

Specific spectral geometric methods were not evaluated or ablated to understand the signal of each individual spectral method. The algorithm also deals with lower sample sizes, though still fails to generalize despite decent training loss. The spectral methods themselves fail on heterogenous tissue, though have been demonstrated in previous work to be useful on homologous structures such as ventral horns, spinal structures, etc. The breadth of the study is limited by the number of algorithms specifically designed for spectral features, and the design of spectral encoders that were designed for phsyics-based tasks, such as solving Partial differential equations. The heat kernel signatures are also predisposed to heavy variance based on the wavelengthe projected and parametrized into the model. It is noted that the validity of geometric spectral methods are not doubted by any means, rather, the objective was to perform a systemwise analysis of performance to analyze the signal carried by spectral features. There is also a limitation as to how the eigenvalues can be obtained, as traditional methods such as schrodinger laplace and traditional rate of change laplace yield heavily invariant solutions. Graphical lapacians were designed for image processing, and capturing spatially differentiable features is difficult to track on a per pixel level, limiting the breadth of information that spectral methods can carry. Nevertheless, these spectral methods are capable of low-data based resolution for tasks that require high inference in data-strained tasks. Simply put, the resolution of these spectral features at the global and local levels are not detectable, and heterogeneous tissue is not readily characterizable by geometric based methods such as fractals, lacunarity, and persistent homology.

## 6 Conclusions

In Conclusion, spectral features provide signal that can boost balanced accuracy, but do not provide enough resolution to be learnable. Spectral features provide a means to outline structures within tissue for models to attend to, but relying on spectral features to train and test a model is limited by the number of algorithms designed to process these features. Geometric methods do however, provide an interpretable means to understand the attention of models within specific regions that is measurable. While fractal based methods provide an interesting method to explore the heterogeneity and patterning of tissue structures, these features on their own are not learnable or attributable when pooled with spectral features. Methods such as fractal convolutions in prior work do prove that fractal are in fact, a hierarchical method of assessing tissue structures. Within Spatial transcriptomics, the resolution spectral mathematical methods is not generalizable across slides, and under performs its counterparts. When predicting within slide spots, the model performs better than traditional methods under sparse conditons. In the future, spectral models can be developed for digital pathology, though under current conditions, spectral features hold great promise as complementary features

## 7 Ethics Statement

All datasets are public and anonymized.

## 8 Author Contributions

All work was performed by the first author, with supervision by the second.

## 11 Data Availability

GBM, BreakHis, and Visium are publically available datasets. All code is available https://github.com/Alejandro21236/SpectralAnalysisPathology

